# A programmable and site-specific anchoring system for biophysical analysis of DNA:protein interactions on large topologically closed DNA molecules

**DOI:** 10.1101/2023.05.12.540605

**Authors:** Neville S. Gilhooly, Stephen C. Kowalczykowski

## Abstract

Single-molecule and bulk biophysical approaches to study protein-DNA interactions on surface-immobilised nucleic acid templates typically rely on modifying the ends of linear DNA molecules to enable surface-DNA attachments. Unless both strands are constrained, this results in topologically-free DNA molecules and the inability to observe supercoiling-dependent biological processes, or requires additional means to micro-manipulate the free DNA end to impose rotational constraints or induce supercoiling. We developed a method using RecA protein to induce the formation of a circularised compliment-stabilised D-loop. The resulting joint molecule is topologically closed, surface anchorable and stable under microfluidic flow. Importantly, the method obviates the need for subsequent manipulation of surface tethered DNA; tethered molecules remain supercoiled and retain accessibility to DNA binding proteins. This approach adds to the toolkit for those studying processes on DNA that require supercoiled DNA templates or topologically constrained systems.

**WHY IT MATTERS:** Supercoiling plays an important role in regulating genetic processes such as DNA replication and transcription. We have developed a facile method to immobilise large (supra diffraction-limited) supercoiled DNA substrates without the need for complex trapping modalities to allow biophysical interrogation of the surface-tethered DNA molecules. This will expand the toolkit for experimentalists interested in studying protein-DNA interactions at the single-molecule or ensemble level requiring the use of topologically closed or supercoiling-dependent systems such as origin-dependent bacterial DNA replication.

## INTRODUCTION

Over the last several decades, biophysical analysis of protein-DNA interactions has, and continues to, shed light on the molecular mechanisms that occur during nucleic acid metabolism including DNA replication, transcription and recombination. Common approaches at both the ensemble and single-molecule level involve the tethering of one or more reaction components to a solid support and measuring the dynamic association of binding partners to the immobilised ligand from bulk solution, yielding thermodynamic and kinetic parameters of the binding process. Examples of these approaches include surface plasmon resonance (SPR) and Co-localisation Single-Molecule Spectroscopy (CoSMoS) (which can also include single-molecule Forster Resonant Energy Transfer (smFRET) analysis) (1, 2). In experiments where the nucleic acid template of interest is tethered to a surface, one can direct the DNA-dependent assembly of complex molecular machines and reconstitute processive biochemical reactions such as DNA replication, transcription, DNA unwinding and DNA end-resection (3-6). Instead of imaging DNA in a random coiled state, microfluidic flow allows one to stretch DNA molecules and resolve individual protein-DNA complexes bound to different loci as well as to track 1D movement of these complexes along the DNA.

Attachment of nucleic acids to surfaces is usually accomplished by modifying the ends of DNA with biotin or digoxygenin moieties to which they can specifically adhere to streptavidin or anti-digoxygenin coated surfaces, respectively. An inevitable consequence of this attachment strategy is that circular DNA species such as plasmids must be linearized prior to end-modification or naturally linear substrates such as Lambda phage DNA are employed. The resulting tethered molecules are not rotationally constrained, i.e., cannot undergo supercoiling. Many processes that occur on DNA *in vivo* are either directly regulated, or affected, by DNA supercoiling. The initiation of bacterial DNA replication at *oriC* shows an absolute requirement for negative supercoiling to allow transient unwinding of the “DNA Unwinding Element” by the initiator protein, DnaA, exposing ssDNA onto which the replicative helicase, DnaB is loaded by DnaC (7, 8). Negative supercoiling also facilitates open complex formation of RNA polymerase at promoters (9). Strategies to impart supercoiling onto tethered linear DNA molecules work by capturing both strands of the free DNA end to a micron-sized bead which can be trapped in a magnetic or optical field. Subsequent rotation of the magnetic field or laser polarization angle results in an applied torque on the DNA which imparts a change in twist and thus supercoiling (10, 11). Importantly, this procedure occurs *in situ* and in the case of magnetic or optical tweezers, relies on complex instrumentation to combine with fluorescence detection. It is therefore clear that it would be advantageous to have a tethering strategy to immobilise supercoiled circular DNA, i.e., intact plasmids so that biological reactions that occur on supercoiled DNA can be interrogated using visual biochemical approaches such as TIRF microscopy, smFRET, etc. without a need for a force imparting modality on the microscope to induce supercoils *in situ*.

A promising approach to make bespoke modifications to plasmid DNA was reported recently that used a large synthetic (18 kbp) plasmid with strategically placed nicking enzyme sites. However, in order to add surface tethering functionality to these molecules they had to be linearised (12). The position of the nicking sites relative to any new inserts into the plasmid would also require subsequent rounds of mutagenesis or cloning. In theory, one could incorporate a biotin modification at the internal nicking site, re-ligate the plasmid, impart negative supercoils with DNA gyrase and isolate via CsCl gradient centrifugation. However, this method is cumbersome requiring specialist molecular biology expertise and results in low yields with non-physiological supercoiling densities. Another approach reported that a pair of bis-PNA openers could be used to melt duplex DNA to which an oligonucleotide could be annealed to form an earring complex i.e., hemi-catenane (13). Synthesis of PNA openers could be prohibitively expensive to some labs, requiring several thousand dollars for each pair which rapidly becomes untenable if one wants multiple choice of anchor location.

We reasoned that we could circumvent these complex manipulations by employing the DNA strand exchange protein, RecA. During the recombinational repair of DNA breaks, RecA protein forms a filament on ssDNA and catalyses invasion of the ssDNA into homologous dsDNA while displacing one of the strands to form a joint molecule intermediate known as a D-loop (14). Including a second ssDNA molecule with sequence complementarity to the displaced strand of the strand exchange reaction D-loop results in a dual-paired, and highly robust, structure termed a complement-stabilized D-loop (cs-D-loop) (15). We report and validate a method that describes the use of RecA and two oligonucleotides that results in a topologically linked site-specific anchor on a large plasmid. We demonstrate the anchoring process on glass coverslips and imaging of single DNA molecules and their *in situ* linearization with a restriction endonuclease. Our approach is easy to implement and requires no sub-cloning to change anchor position – it is programmable by choice of homology region. The method should be of general interest to those wishing to study biological processes on immobilised negatively supercoiled DNA molecules and will allow processes such as origin-dependent bacterial replication to be reconstituted at the single-molecule level.

## MATERIALS AND METHODS

### Construction of a large OriC-containing plasmid

A region of the *E. coli* (MG1655) genome containing the *rrnC* operon was amplified by PCR (Phusion, NEB) using the following primers:

rrnC_for_NotI-GGAGACGCGGGCCGCTGAAAAAATGCGCGGTCAG) and rrnC_rev_BamHI-GGAGACGGATCCTATTTACTTCGGAGAGGGTTATTTCAG. PCR products were digested with BamHI and NotI (NEB) and ligated into BamHI/NotI-digested pACYC-Duet-1 vector (Novagen). The resulting ∼9.6 kbp plasmid was then cut with BamHI and ligated to a ∼9.5 kbp BamHI-digested PCR product from a region of *E. coli* (MG1655) genomic DNA. Synthetic DNA fragments containing an OriC (TTGTCGACGGAGCTCGAATTCGGATGGATCCTGGGTATTAAAAAGAAGATCTATTTATTTAGAGATCTGTTCTATTGTGATCTCTTATTAGGATCGCACTGCCCTGTGGATAACAAGGATCCGGCTTTTAAGATCAACAACCTGGAAAGGATCATTAACTGTGAATGATCGGTGATCCTGGACCGTATAAGCTGGGATCAGAATGAGGGGTTATACACAACTCAAAAACTGAACAACAGTTGTTCTTTGGATAACTACCGGTTGATCCAAGCTTCCTGACAGAGTTATCCACAGTAGATCGCACGATCTGTATACTTATTTGAGTAAATTAACCCACGATCCCAGCCATTCTTCTGCCGGATCTTCCGGAATGTCGTGATCAAGAATGTTGATCTTCAGTGTTTCGCCTGTCTGTTTTGCACCGGAATTTTTGAGTTCTGCCTCGAGTTTATCGATAGCCCCACAAAAGGTGTCATATTCACGACTGCCAATACCGATTGCGCCAAAGCGGACTGCAGAAAGATCGGGCTTCTGTTCCTGCAATGCTTCATAGAAAGGAGAAAGGTTGTCCGGAATATCTCCGGCACCGTGGGTGGAGCTGATAACCAGCCAGATCCCTGAGGCAGGTAAATCTTCTAACAGCGGACCGTGCAGCGTTTCGGTGGTAAAACCCGCCTCTTCCAGCTTTTCAGCCAGGTGTTCTGCTACATATTCGGCACCGCCGAGGGTGCTGCCGCTGATAAGAGTGATATCTGCCATCCAGGGTGGTTTTTCTTTTCACCAG) and F1-origin (GGGGAGAGGCGGTTTGCGTATTGGGTAAGAATTAATTCATGAGCGGATACATATTTGAATGTATTTAGAAAAATAAACAAATAGGGGTTCCGCGCACATTTCCCCGAAAAGTGCCACCTGAAATTGTAAACGTTAATATTTTGTTAAAATTCGCGTTAAATTTTTGTTAAATCAGCTCATTTTTTAACCAATAGGCCGAAATCGGCAAAATCCCTTATAAATCAAAAGAATAGACCGAGATAGGGTTGAGTGTTGTTCCAGTTTGGAACAAGAGTCCACTATTAAAGAACGTGGACTCCAACGTCAAAGGGCGAAAAACCGTCTATCAGGGCGATGGCCCACTACGTGAACCATCACCCTAATCAAGTTTTTTGGGGTCGAGGTGCCGTAAAGCACTAAATCGGAACCCTAAAGGGAGCCCCCGATTTAGAGCTTGACGGGGAAAGCCGGCGAACGTGGCGAGAAAGGAAGGGAAGAAAGCGAAAGGAGCGGGCGCTAGGGCGCTGGCAAGTGTAGCGGTCACGCTGCGCGTAACCACCACACCCGCCGCGCTTAATGCGCCGCTACAGGGCGCGTCCCATTCGCCAATCCGGATATAGTTCCTCCTTTCAGCAAAAAACCCCTCAAGACCCGTTTAGAGGCCCCAAGGGGTTATGCTAGTTATTGCTCAGCGGTGGCAGCAGCCAACTCAGCTTCCTTTCGGGCTTTGTTAGCAGCCGGATCTCAGTGGTGGTGGTGGTGGTGCTCGAGTGCGGCCGCAAGCTTGTCGACGGAGCTCGAATTCGGAT) sequence (Genewiz) were then introduced via Gibson assembly into the NarI-digested vector resulting in a final 20,359 bp plasmid harbouring OriC and the *rrnC* operon transcription cassette denoted as pNGho.

### Plasmid DNA purification and oligonucleotides

pNGho was propagated in a 1 L culture of DH5α and grown at 30°C to minimise recombination events. Large scale plasmid purification was accomplished using a Qiagen maxi-prep kit followed by clean-up of the DNA by 4-20% sucrose gradient centrifugation in 10 mM Tris and 1 mM EDTA (TE, pH 7.5) buffer + 1 M NaCl. DNA was ethanol precipitated and resuspended in TE buffer, aliquoted and stored at -20°C until use.

Oligonucleotides used in D-loop formation assays were synthesised and HPLC purified by IDT and sequences shown below:

<u>70-mer oligo1:</u>

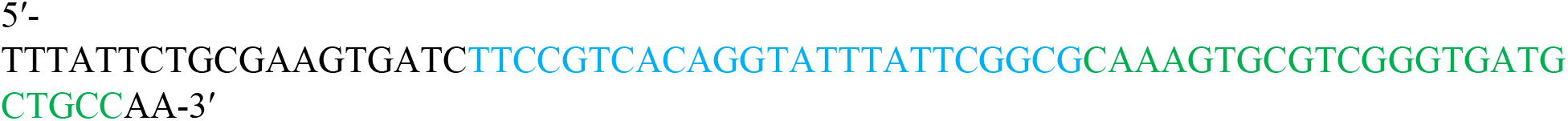

<u>106-mer complement oligo:</u>

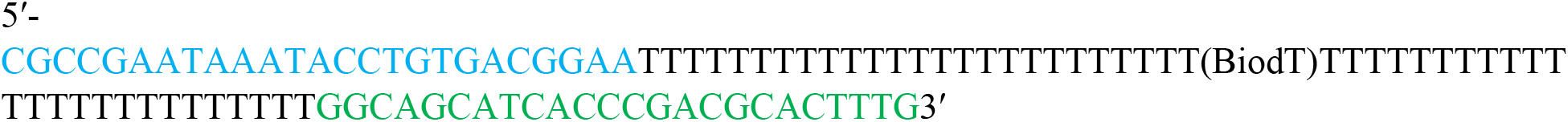

The regions in the compliment oligo that base pair with the displaced strand of the D-loop formed with oligo1 are shown in blue and green, respectively. Note the compliment oligo contains an internal biotin-dT modification.

### D-loop assays

Oligonucleotides used as substrates in D-loop assays were radiolabelled at their 5′-end using T4 Polynucleotide Kinase (T4 PNK - NEB) and adenosine triphosphate [γ-^32^P] (Perkin Elmer) following the manufacturer’s instructions. Following heat inactivation of PNK, unincorporated nucleotides were removed via passage through a Microspin G-25 column (Cytiva). RecA was purified as described in (16). RecA nucleoprotein filaments were formed on ssDNA oligonucleotides (concentration denoted in figure legends) at a ratio of 1 RecA monomer per 2 nucleotides in a buffer containing Tris(OAc) (25 mM, pH 7.5), Mg(OAc)_2_ (10 mM), DTT (1 mM) and adenosine 5′-[γ-thio]triphosphate (1 mM) for 10 mins at 37°C. Joint molecule formation was initiated by the addition of supercoiled plasmid DNA (14.5 nM, molecules). The reaction was allowed to proceed for 30 mins before the addition of an equal volume of stop buffer (Proteinase K (1 mg.mL^-1^), EDTA (100 mM), Ficoll 400 (10%), SDS (5%), bromophenol blue (0.125%), and xylene cyanol (0.125%). Samples were deproteinised for 30 mins at room temperature prior to electrophoresis through a native 1% agarose gel, formed in 40 mM Tris-base, 1 mM EDTA and 0.114% acetic acid, and run at 50V for 3 hours. Gels were dried onto DEAE paper and visualised using a Storm phosphorimager. The percentage of D-loops formed was calculated using ImageQuant software (Cytiva) by expressing the yield relative to the amount of input plasmid (14.5 nM, molecules).

### Purification of plasmids with a circularised D-loop

A preparative scale (267 μL) of the optimised D-loop reaction was accomplished by incubating the targeting 70-mer oligo (200 nM, molecules) with RecA (28 μM) for 10 minutes in a reaction buffer consisting of Tris(OAc) (25 mM, pH 7.5), Mg(OAc)_2_ (10 mM), DTT (1 mM) and ATPγS (1 mM). Supercoiled plasmids (14.5 nM) were added to initiate D-loop formation and the reaction was allowed to continue for 30 minutes. The 106-mer biotin-modified oligo was then added to a final concentration of 200 nM, molecules and complement stabilised D-loops allowed to form for 10 minutes. The mixture was extracted with an equal volume of a 50:50 mixture of phenol and chloroform and the aqueous phase desalted using a Biospin 6 column. The solution was supplemented with 0.1 vol of 10X T4 ligase buffer (NEB) and T4 ligase (30 U.μL^-1^, NEB) and incubated at 25°C for 2 hours. DNA was ethanol precipitated and resuspended in TE and stored at 4°C until use.

### Flow-cell fabrication and coverslip functionalisation

Flow-cells and coverslips were constructed and treated as described in (17) but with the following modifications: instead of treating coverslips with 3-APTES and functionalisation with mPEG-NHS ester, coverslips were functionalised by applying a 50 μL droplet of a 200 mg/mL total mixture of 95% mPEG silane: 5% biotin-mPEG silane (MW 5000 each, Laysan bio) dissolved in 90% EtOH + 10% acetic acid and incubating at 50-60°C for 16 hours in the dark. Coverslips with rinsed thoroughly with Milli-Q water, dried with nitrogen gas and stored in Parafilm-wrapped Falcon tubes at -20°C until use. Prior to experiments, flow-cell surfaces were further functionalised at room temperature by injecting 100 μL of reaction buffer (see DNA tethering and single-molecule fluorescence imaging section) supplemented with neutravidin (0.1 mg.mL^-1^) and incubating for 5 minutes. The flow-cell was washed with 100 μL of reaction buffer and then further passivated by incubation with 100 μL of reaction buffer supplemented with Roche Blocking Reagent (150 μg.mL^-1^) for 15-30 minutes. The flow-cell was then washed with 100 μL of reaction buffer prior to experiments.

### DNA tethering and single-molecule fluorescence imaging

Anchoring of DNA molecules was performed in a de-gassed buffer (typically used to support reconstituted DNA replication reactions) containing HEPES-KOH (50 mM, pH 8.0), Mg(OAc)_2_, (10 mM), DTT (50 mM), sucrose (15%), dATP, dTTP, dCTP, dGTP (40 μM each), GTP, CTP, UTP (200 μM), and ATP (1 mM), SYTOX green (Invitrogen) (50 nM). To linearize anchored plasmids, BmtI-HF was introduced into the flow-cell at 2 U.μL.^-1^. Experiments were performed at 37°C using an Eclipse TE2000-U, inverted TIRF microscope (Nikon) as described in (17). SYTOX Green was illuminated continuously with 488 nm excitation at a frame rate of 20 s^-1^ with an illumination power of 60 μW (measured at the objective). Prior to data analysis, movies were median averaged over 5 frames and background subtracted using a rolling ball size of 40 pixels (Fiji (18)). Anchored plasmid molecules were detected using the ThunderSTORM V.13 plugin in Fiji (18, 19) and molecule lengths are reported as full width at half maximum (FWHM) derived from the fitted values of the standard deviation from a 2D Gaussian distribution. Molecule lengths in the presence of microfluidic flow were determined by manually measuring the end-to-end distance of DNA molecules in Fiji (18).

## RESULTS

We sought to create a method to allow the anchoring of topologically closed DNA molecules to surfaces. The key attributes of this method were that it should (1) be cost-effective, (2) not require laborious cloning steps to change the anchor position, (3) not require the use of nicking endonucleases, and (4) work on molecules large enough for fluorescence imaging, i.e., capable of spatially resolving protein-DNA complexes when molecules are extended in microfluidic flow. We reasoned that RecA could replace the function of expensive bis-PNA openers, used previously to assemble “DNA earrings” (13), by providing the essential locus-specific strand pairing activity via the formation of a D-loop (Figure 1A). To that end, we constructed a ∼20 kbp plasmid that we used to site-specifically attach to a glass surface for single-molecule imaging while retaining its negatively supercoiled state. To do this, we optimised the D-loop formation assay to maximise the yield of joint molecules by titrating RecA nucleoprotein filaments assembled on a 70-mer oligonucleotide against a fixed plasmid concentration (Figure 1B). ATPγS was used to prevent filament turnover and product dissociation. We found the approach to saturation of D-loop yield occurred at ∼100 nM of the input D-loop forming oligonucleotide, and D-loop formation absolutely required the presence of RecA protein (Figure 1C). We subsequently used the D-loop forming oligonucleotide at 200 nM to ensure maximal yields.

**Figure 1.**
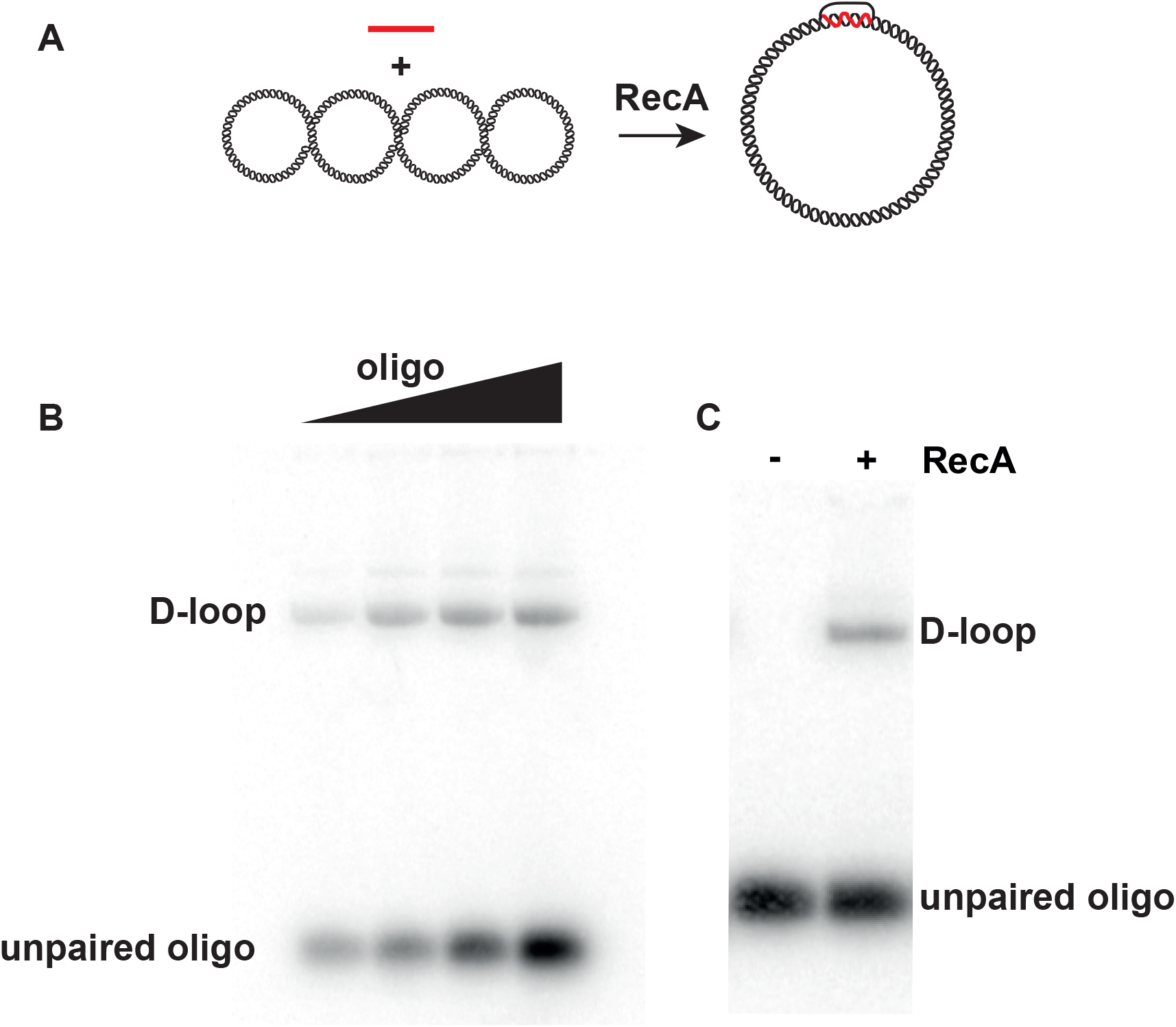
Optimisation of RecA-dependent D-loop formation. (A) Schematic of a reaction between a negatively supercoiled plasmid DNA and oligonucleotide with homologous sequence to the plasmid. RecA catalyses DNA strand exchange between the oligonucleotide and plasmid inducing the formation of a D-loop structure. (B) Titration of RecA presynaptic filaments (10, 20, 50 and 100 nM molecules – RecA:ssDNA = 2:1) against a fixed concentration of plasmid (14.5 nM, molecules). (C) RecA-dependency of D-loop formation. D-loops were separated from the free oligo by native agarose gel electrophoresis.

We next developed a protocol whereby an oligonucleotide containing an internal biotin modification and 5′-phosphate was annealed to the displaced strand in such a way that its 5′- and 3′-paired ends are juxtaposed for ligation. Subsequent ligation would then yield a circularised oligonucleotide base-paired to the displaced strand of the D-loop catalysed by RecA - in other words, a topologically entrapped linkage on a circular DNA molecule (Figure 2A). In this experiment, the second 106-mer oligonucleotide containing the internal biotin modification is radiolabelled and the 70-mer used previously is cold. We used heat treatment after ligation as an indication of successful topological entrapment of the 106-mer on DNA. In the absence of ligase, all D-loop products were heat-labile (Figure 2B - lane 2). However, in the presence of ligase, a substantial portion of the products were heat-stable indicating topological linkage between the plasmid and labelled oligo (Figure 2B – lane 3, red asterisk). A product that runs near the wells of the gel is also observed after heat treatment (Figure 2B - lane 2) which we interpret as cs-D-loop containing plasmids that spuriously and *in trans* reannealed prior to, or during, electrophoresis. The biotin-containing oligonucleotide itself does not promote D-loop formation in the presence of RecA (Figure 2C – lane 2), probably due to the inverted and non-contiguous homologous regions, which are incompatible with known biophysical properties of RecA:ssDNA filaments (20). However, DNA annealing occurs when RecA filaments are assembled first on the cold 70-mer to form a D-loop, indicating formation of a cs-D-loop (Figure 2B – lane 3). We found that the order of addition of each oligo did not impact cs-D-loop formation, however there was an absolute dependency of the 70-mer oligonucleotide. We interpret this to mean that D-loop formation with the 70-mer precedes annealing of the 106-mer which is not inhibited by the presence of free RecA protein in the reaction. Taken together, the dependency of the 70-mer and ligase activity to observe gel-stable and heat-stable products, respectively, indicates the formation of a circularised cs-D-loop.

**Figure 2.**
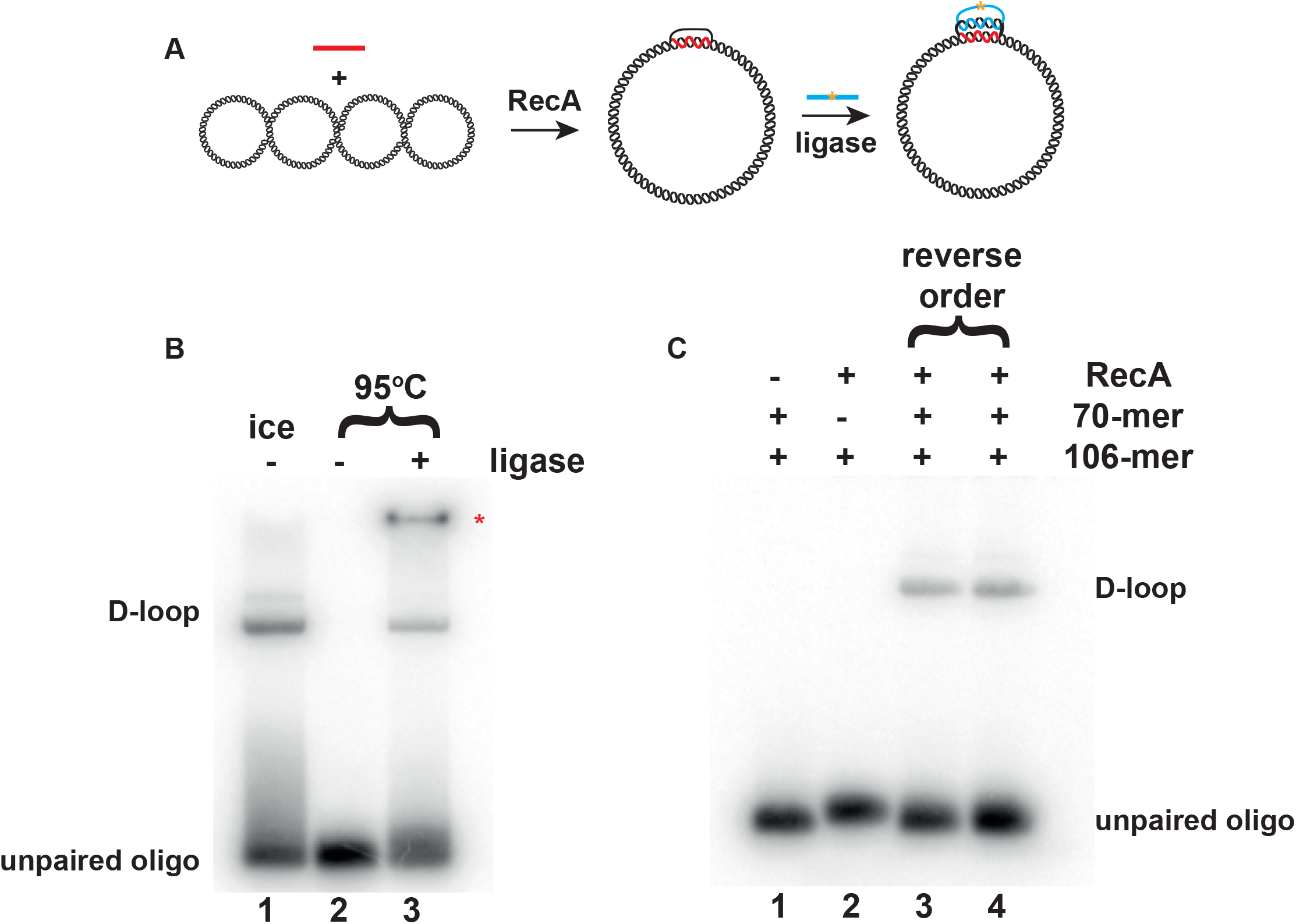
Production of a topologically closed cs-D-loop. (A) Schematic of a reaction that generates a cs-D-loop with juxtaposed ligatable ends. After RecA induces the formation of a D-loop using an oligonucleotide with sequence homology to the negatively supercoiled plasmid, a second oligonucleotide that is complimentary to the displaced strand can anneal forming a cs-D-loop that is topologically trapped upon ligation. A gold asterisk indicates the position of the biotin-dT moiety. (B) Verification of cyclization of the cs-D-loop using heat-stability as an indirect readout of topological entrapment. Red asterisk indicates a spurious annealing product after heat denaturation (C) Oligo and RecA-dependencies of cs-D-loop formation.

We further confirmed that these plasmids, by virtue of their biotinylated hemi-catenane, could be anchored to glass surfaces and imaged with TIRF microscopy (Figure 3). Anchored plasmids were visualised by using the intercalating dye, SYTOX Green. In the absence of microfluidic flow, plasmids appear as bright foci due to the plasmids adopting a random coil confirmation constrained around the anchor point with apparent FWHM of 549 ± 99 nm (Figures 3A and 3C). The airy disc FWHM of our optical system would be ∼180 nm based on the emission properties of SYTOX Green, indicating that while these plasmids appear as circular foci, they occupy a size larger than the point spread function of our microscope, consistent with their supra-diffraction limit size. In the presence of flow, the plasmids can be further extended 3.6-fold from their anchor points and now resemble linear stretched DNA molecules of mean length 2.0 ± 0.2 μm. We could also linearize these anchored plasmids *in situ* indicating that the attachment process did not affect the accessibility to DNA binding proteins and, importantly, that the hemi-catenane structure was stable under flow, allowing hydrodynamic extension of the linearized plasmid molecules (Figures 3B & C). Linear molecules had an apparent length of 5.2 μm ± 0.35 μm, more than double the size of the supercoiled circular molecules, as expected. The stability of the attachment was confirmed by cycling the flow on and off following linearization of the plasmid, indicating that the anchor survives the sudden changes of force exerted by plasmid linearization as well as pulsatile flow (Figure 3B).

**Figure 3.**
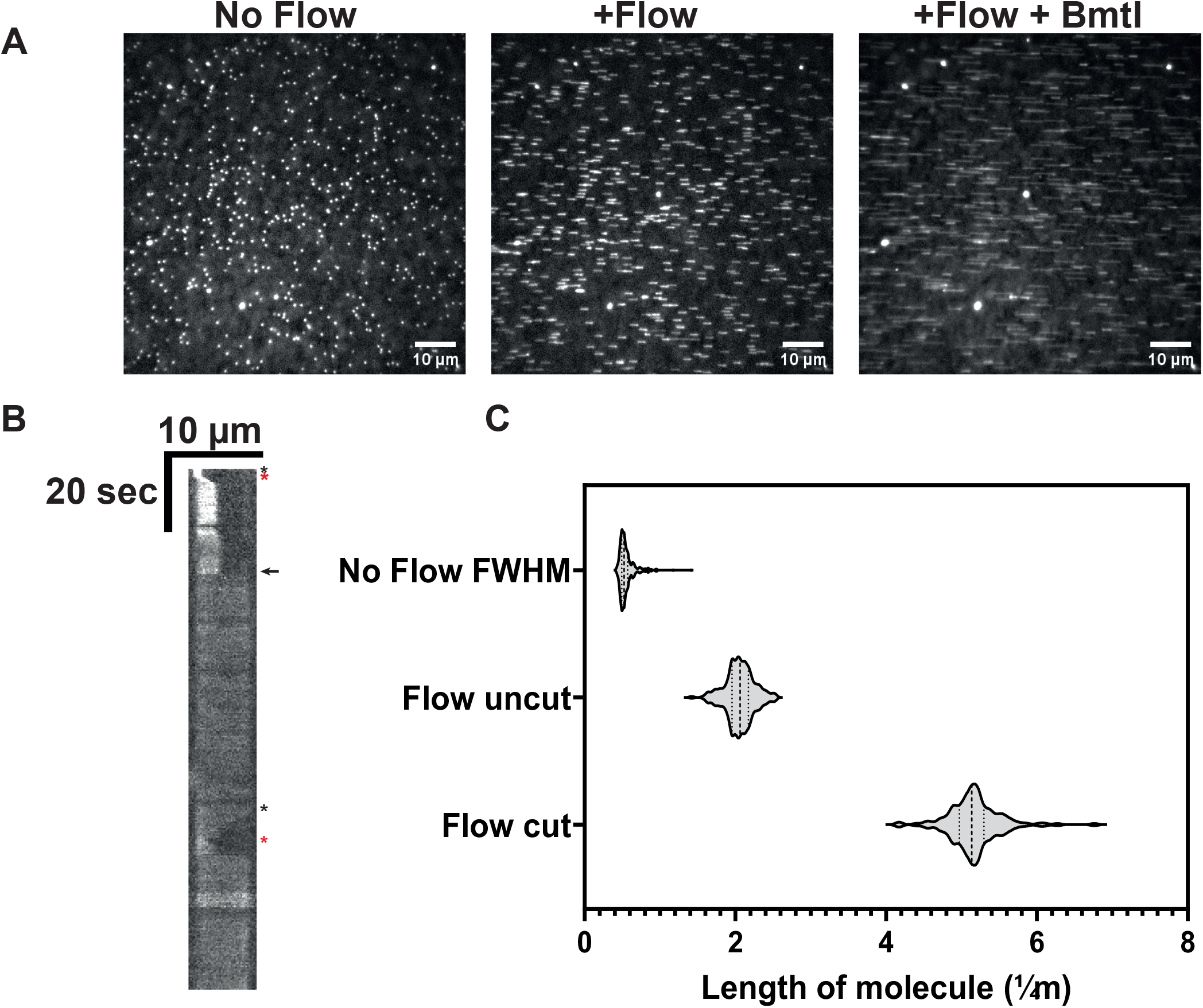
Single-molecule imaging of anchored plasmids and their *in situ* linearization with a restriction enzyme. (A) (*left*) Micrograph of anchored plasmids stained with SYTOX Green in the absence of microfluidic flow. (*center*) The same field of view experiencing microfluidic flow. (*right*) The same field of view under flow after cleavage with BmtI-HF restriction endonuclease. (B) Kymograph of an anchored DNA molecule. The black arrow indicates when the molecule was cleaved by BmtI-HF; note the sudden extension of the DNA. Black and red asterisks indicate when flow was turned off and on, respectively. (C) Quantification of molecule lengths in the presence and absence of flow and after linearization, respectively.

## DISCUSSION

We describe a straightforward “one pot” reaction to immobilise large plasmid molecules onto solid-phase supports using biotin-streptavidin conjugation chemistry that simply requires RecA protein and two oligonucleotides, one with an internal biotin modification. Assuming an average negative superhelical density (σ) *in vivo* of -0.06 (21), the plasmid has a Linking number of 1841 comprising a writhe of -117 in order to maintain the initial Twist of 1958. Formation of the D-loop will impart a change in Linking number (ΔLk) of 6.6. Recalling that σ = ΔLk/Lk_0,_ the anchor does not substantially change the net negative supercoiled nature of the plasmid (σ_final_ = -0.056). Importantly, this means our method can be used to specifically anchor negatively supercoiled plasmids allowing biophysical integration of processes that occur on, or require, supercoiled DNA templates.

While we used glass surfaces for TIRF microscopy experiments, any streptavidin-coated surface could be applicable, such as SPR chips for ensemble biophysical measurements of protein-DNA interactions on supercoiled DNA. An important feature of the method is that the anchor survives release of superhelical energy and subsequent flow extension of the DNA. This will permit studies that require assembly of protein complexes on supercoiled DNA and subsequent visualisation of optically-resolved (due to the ∼5 μm template length) protein complexes along the *in situ* linearized DNA molecule. This method could be also be applied to smFRET or tethered particle motion-type experiments to observe the association of proteins on supercoiled DNA. Our method bypasses the need for PNA synthesis which can cost thousands of dollars. It also doesn’t require magnetic or optical trapping modules to induce supercoiling *in situ* which has an additional benefit of not requiring complicated dual modality instruments such as hybrid magnetic tweezer instruments to perform visual biochemical experiments on supercoiled templates.

Because we employ a homology-directed method, the anchor position can be easily relocated by changing the sequence of the D-loop forming oligonucleotide bound by RecA instead of having to perform further rounds of cloning to the starting plasmid. RecA protein is also commercially available from NEB. Therefore, we think this method is both generalisable and accessible to many biophysics labs and it doesn’t require complex molecular biology or protein-purification expertise.

## Acknowledgments

These studies were supported by an Anna D. Barker Basic Cancer Research Fellowship (AACR) to N.S.G. and NIH R35 GM131900 to S.C.K.

